# Defensin-driven viral evolution

**DOI:** 10.1101/2020.05.08.079574

**Authors:** Karina Diaz, Ciara T. Hu, Youngmee Sul, Beth A. Bromme, Nicolle D. Myers, Ksenia V. Skorohodova, Anshu P. Gounder, Jason G. Smith

**Author notes:** Address correspondence to Jason G. Smith.

## Abstract

Enteric alpha-defensins are potent effectors of innate immunity that are abundantly expressed in the small intestine. Certain enteric bacteria and viruses are resistant to defensins and even appropriate them to enhance infection, despite neutralization of closely related microbes. We therefore hypothesized that defensins impose selective pressure during fecal-oral transmission. Upon passaging a defensin-sensitive serotype of adenovirus in the presence of a human defensin, mutations in the major capsid protein hexon accumulated. In contrast, prior studies identified the vertex proteins as important determinants of defensin antiviral activity. Through infection and biochemical assays, we found that although all major capsid proteins serve a critical role in defensin-mediated neutralization, hexon is the sole determinant of enhancement. These results extensively revise our understanding of the interplay between defensins and non-enveloped viruses. Furthermore, they provide a feasible rationale for defensins shaping viral evolution, resulting in differences in infection phenotypes of closely related viruses.

**Author Summary:** Defensins are potent antimicrobial peptides that are found on human mucosal surfaces and can directly neutralize viruses. They are abundant in the small intestine, which is constantly challenged by ingested viral pathogens. Interestingly, non-enveloped viruses, such as adenovirus, that infect the gastrointestinal system are unaffected by defensins or can even appropriate defensins to enhance their infection. In contrast, respiratory adenoviruses are neutralized by the same defensins. How enteric viruses overcome defensin neutralization is not well understood. Our studies are the first to show that defensins can drive the evolution of non-enveloped viruses. Furthermore, we identify important components within human adenovirus that dictate sensitivity to defensins. This refined understanding of defensin-virus interactions informs the development of defensin-based therapeutics.

## Introduction

Defensins are small, cationic, and amphipathic peptides that constitute a conserved component of the innate immune system [1, 2]. Expression of these host defense peptides by humans, despite the presence of a refined adaptive immune system, highlights their key role in protection from microbes. Although there are examples of both α- and β-defensins with antibacterial activity and antiviral activity against enveloped viruses, only α-defensins affect infectivity of non-enveloped viruses [1, 3, 4]. There are two major classes of α-defensins: myeloid, which are produced by neutrophils, and enteric, which are secreted in the genitourinary tract and by Paneth cells in the crypts of the small intestine. Human defensin 5 (HD5) is the most abundantly expressed human enteric α-defensin and has the greatest inhibitory activity against human non-enveloped viruses, including polyomavirus, papillomavirus, and some serotypes of adenovirus (AdV) [5-12]. Although differing in some respects between viruses, a conserved mechanism by which non-enveloped viruses are neutralized by α-defensins has emerged. In essence, stabilization of the capsid by α-defensin binding leads to changes in uncoating and intracellular trafficking, thereby preventing the genome from reaching the nucleus to initiate replication [1, 5, 6, 8-11, 13-15].

Despite their broad antimicrobial activity, enteric α-defensins are not able to inhibit all non-enveloped viruses. Echovirus, reovirus, and enteric AdVs from both humans and mice are resistant to enteric α-defensin inhibition [9, 16-18]. One hypothesis to explain these observations is that resistance stems from evolutionary pressure imposed by enteric α-defensins during fecal-oral transmission. Consistent with this hypothesis, rather than kill the enteric bacterial pathogen shigella, HD5 was recently found to promote its cell binding and infection [19, 20]. And, enteric α-defensins play a role in shaping the microbial communities of the gastrointestinal tract through differential susceptibility of commensal bacteria to defensin killing [21-23]. Collectively, these observations suggest that defensin-driven evolution of enteric microbes is a common cross-kingdom occurrence.

To directly test the ability of enteric α-defensins to drive viral evolution, we used human AdV (HAdV). We previously demonstrated that HAdV infection can either be neutralized, resistant to, or enhanced by HD5, depending on serotype [9]. Thus, the naturally occurring diversity of HAdVs is an appealing substrate to identify viral determinants for neutralization and enhancement by defensins. The icosahedral AdV capsid consists of three major proteins: hexon, penton base (PB), and fiber. In previous studies, we used a rational design approach to identify the vertex proteins, fiber and PB, as determinants of HD5 neutralization [9]. Here, we evolved HD5 resistance in a defensin-sensitive serotype. From this, we identified a hypervariable loop in hexon as a novel determinant that both proves the capability of enteric α-defensins to drive viral evolution and substantially revises our understanding of the mechanism of HD5 neutralization.

## Results

### Hexon hypervariable region 1 is a novel determinant of HD5 sensitivity

To determine whether HD5 could impose a selective force for HAdV evolution, we utilized a previously described HAdV-5-based “mutator” vector encoding a polymerase with reduced fidelity to facilitate *de novo* mutagenesis [24]. Over 50 passages, we found that the HD5 IC_90_ used for selection increased 7-fold from ∼2.5 µM to ∼17.5 µM (Fig. 1A). We expanded the viral pool from every 5^th^ round of HD5 selection and the 10^th^ and 20^th^ rounds of passaging control virus. As expected, there was a significant positive correlation between the round of selection and the HD5 IC_50_ of the population (Fig. 1C). Whole genome sequencing yielded an average of 6000 mapped reads per base [25]. Both the initial inoculum and the passaged control samples contained only low frequency (<1%) mutations. In contrast, we observed numerous mutations across the viral genome that exceeded 1% frequency in the HD5 selected samples (Table 1). Mutations in the DNA binding protein (V340A), protein VI (A225T), protein VII (G50D), 52K protein (H193Y), and IIIa (A39) reached a frequency of >50%; however, none of these mutations stayed above this threshold (Fig. 1B). In contrast, mutations in polymerase (R149H, P132, and C228), the U Exon (G5D), hexon (I114K and E154K), and L4-100K (G745D) became fixed in the population as early as the 15^th^ round of selection (Figs. 1A and 1B). In addition, a third mutation in hexon (E424K) was trending towards fixation (Fig. 1A).

**Table 1:**
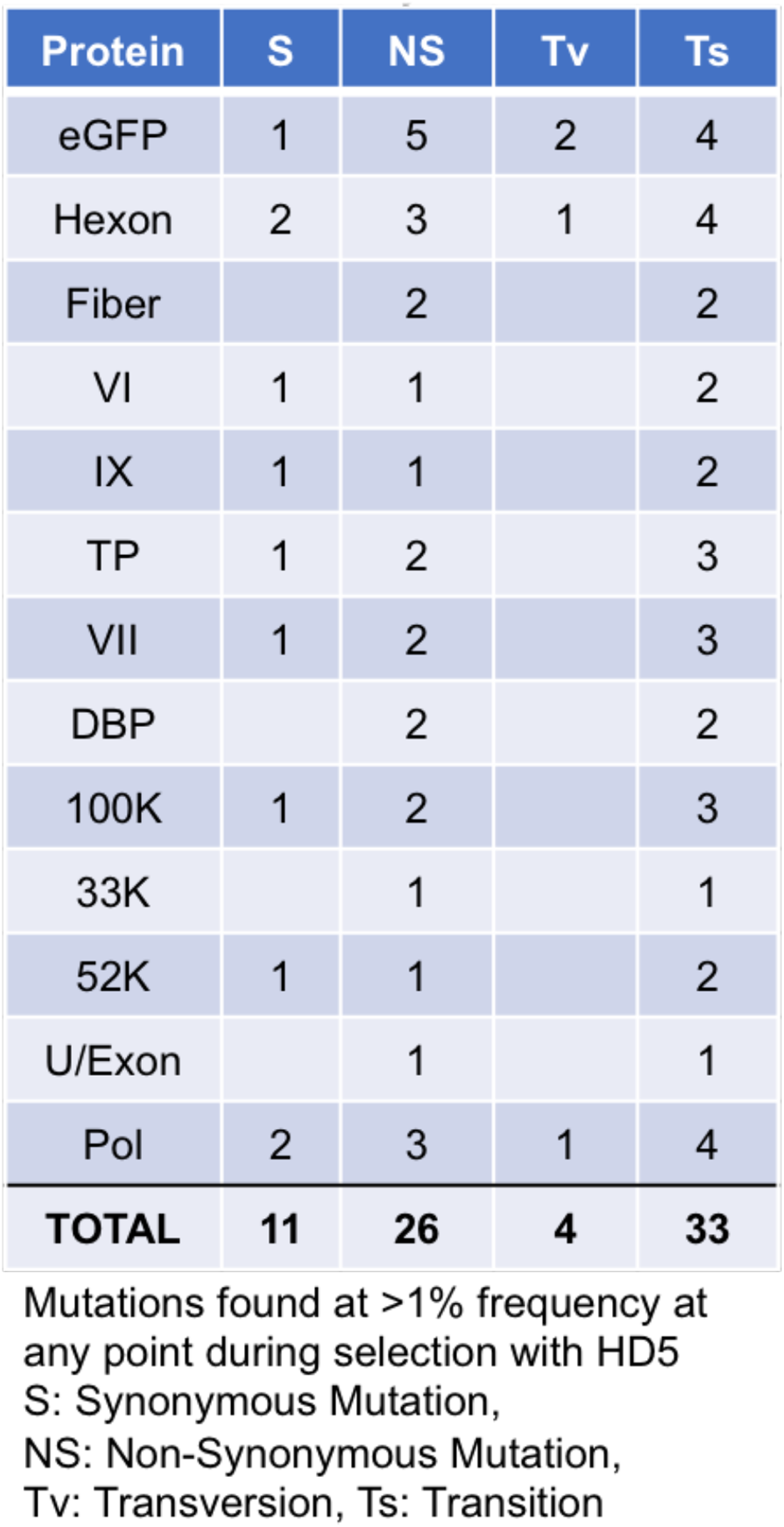
Summary of Mutations

| Protein | S | NS | Tv | Ts |
| --- | --- | --- | --- | --- |
| eGFP | 1 | 5 | 2 | 4 |
| Hexon | 2 | 3 | 1 | 4 |
| Fiber |  | 2 |  | 2 |
| VI | 1 | 1 |  | 2 |
| IX | 1 | 1 |  | 2 |
| TP | 1 | 2 |  | 3 |
| VII | 1 | 2 |  | 3 |
| DBP |  | 2 |  | 2 |
| 100K | 1 | 2 |  | 3 |
| 33K |  | 1 |  | 1 |
| 52K | 1 | 1 |  | 2 |
| U/Exon |  | 1 |  | 1 |
| Pol | 2 | 3 | 1 | 4 |
| **TOTAL** | **11** | **26** | **4** | **33** |

Mutations found at >1% frequency at any point during selection with HD5 S: Synonymous Mutation, NS: Non-Synonymous Mutation, Tv: Transversion, Ts: Transition

**Figure 1.**
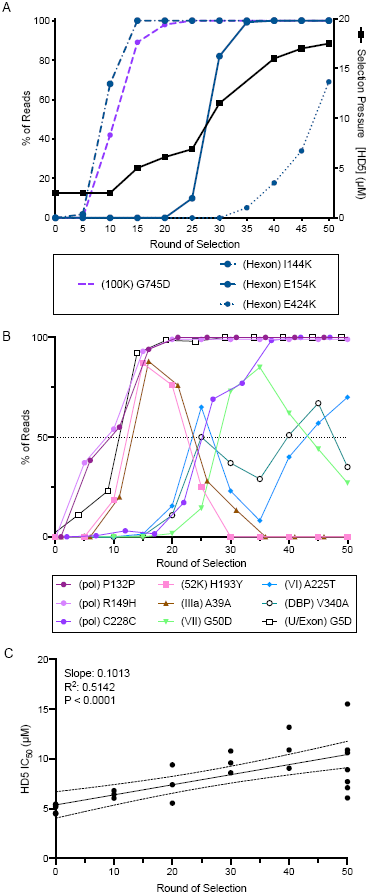
Selection of HD5-resistant HAdV-5. HAdV-5 “mutator” virus was pre-bound to 293β5 cells and then incubated with HD5 to select for resistant viruses. (A) The right y-axis indicates the concentration of HD5 used for selection (solid black line). The left y-axis indicates the percentage of reads containing the denoted mutations in hexon (blue) and L4-100K (purple) in pools of virus expanded from the bulk selected population. (B) All mutations in other proteins that were found in at least 50% of the population during selection. (C) HD5 IC_50_s of pools of virus expanded from the bulk selected population was determined on 293β5 cells. Each data point is an independent experiment, and linear regression with 95% confidence bands is graphed.

Each of the hexon mutations introduced positive charge in one of two hypervariable regions (HVRs), HVR1 (I144K and E154K) or HVR7 (E424K), located on the outer face of hexon (Fig. 2C); therefore, we focused on their contribution to HD5 resistance. We plaque purified viruses with one, two, or three of these hexon mutations in combination with the L4-100K mutation. We were unable to isolate viruses that contained only hexon mutations in the absence of the L4-100K mutation, perhaps due to the role of this protein as a chaperone for the proper folding, trimerization, and transport of hexon [26, 27]. Thus, we engineered a virus containing both HVR1 mutations in the absence of the L4-100K mutation. Viruses that contained only the mutation in L4-100K had an IC_50_ equivalent to that of HAdV-5 (Fig. 2A, salmon). Viruses with the L4-100K mutation and only one (pink) or both (blue) of the hexon mutations in HVR1 had a significantly higher IC_50_ compared to the starting population but were equivalent to each other. The presence of all three hexon mutations (green) resulted in a ∼3-fold increase in IC_50_ from the starting population. Interestingly, the virus engineered with the two mutations in HVR1 but without the mutation in L4-100K (purple) was just as resistant to HD5 as viruses with all three hexon mutations and L4-100K. Taken together, these results indicate an important role for HVR1 and HVR7 in sensitivity of HAdV-5 to neutralization by HD5.

**Figure 2.**
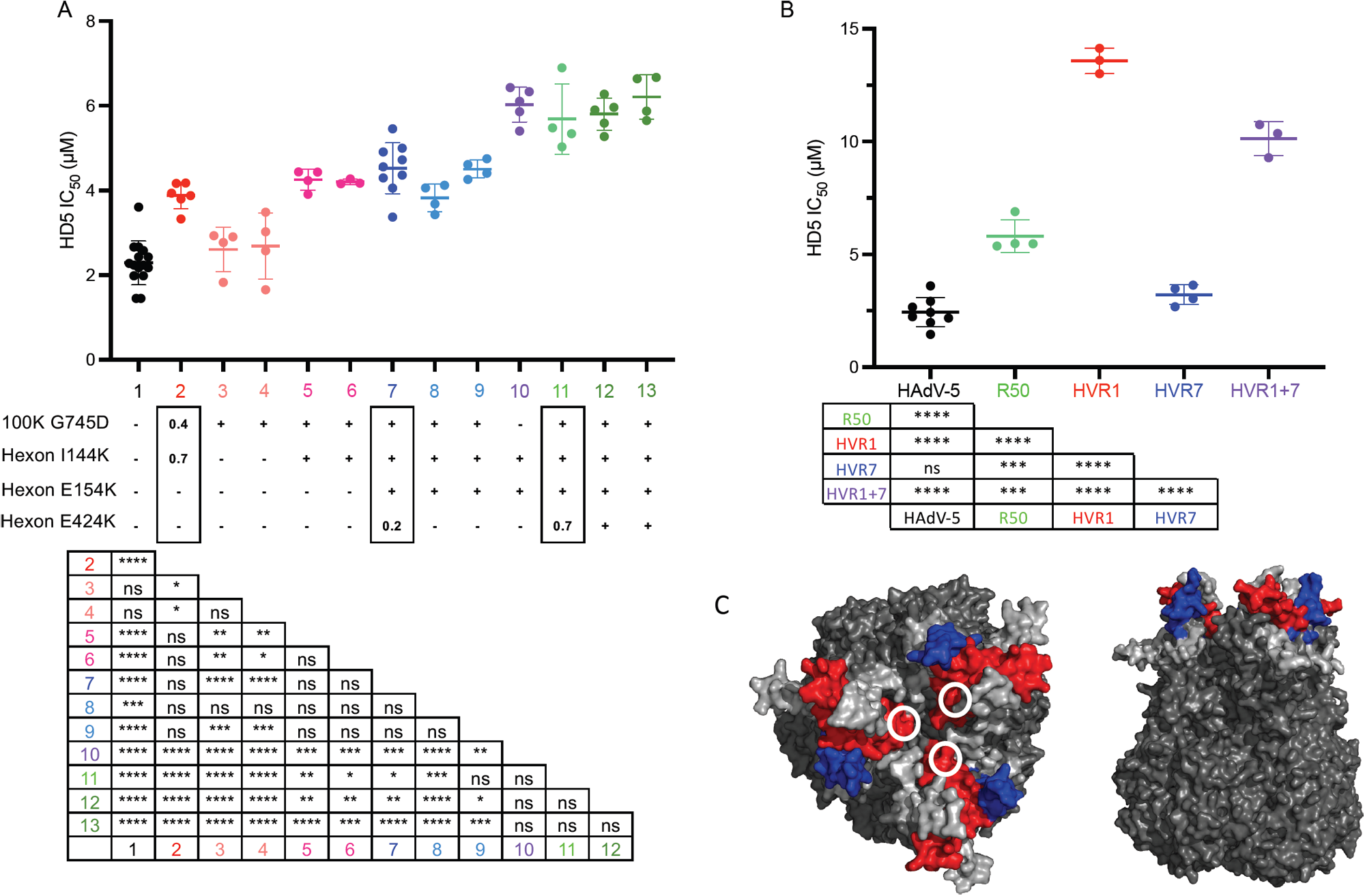
Hexon HVR1 is a determinant of HD5-mediated neutralization. (A) The HD5 IC_50_s of HAdV-5 “mutator” virus (column 1, black); pools of virus expanded from rounds 10 (column 2, red), 40 (column 7, dark blue), and 50 (column 11, light green); plaque purified viruses from rounds 10 (columns 3-4, salmon; columns 5-6, pinks), 40 (columns 8 and 9, blues), and 50 (columns 12 and 13, greens) of the selection; and an engineered virus containing only the indicated hexon mutations (column 10, purple) were determined on 293β5 cells. The presence of each hexon and L4-100K mutation is denoted below the graph. For the pooled viruses, the fraction of reads containing each mutation is indicated within the rectangles. Each data point is an independent experiment, and bars are the mean ± SD. The results of one-way ANOVA with Tukey’s multiple comparison test are indicated by asterisks below. (B) The HD5 IC_50_s of the indicated viruses were determined on A549 cells. Each data point is an independent experiment, and bars are the mean ± SD. Virus expanded from the 50^th^ round of selection (R50) is equivalent to sample 11 in panel A. Results of one-way ANOVA with Tukey’s multiple comparisons test are shown by asterisks below. (C) Space-filling model of the structure of a HAdV-5 hexon trimer (PDBID:6CGV [36]) from top and side views. HVR1 (blue) is modeled using the structure of the shorter HVR1 from species D HAdV-26 (PDBID:5TX1 [37]). White circles indicate the position of E424 in HVR7 (red). All other HVRs are light gray.

To further explore the potential importance of HVR1 and HVR7 for HD5 interactions, we created chimeric viruses at these regions between HAdV-5 and HAdV-64, an HD5-enhanced serotype [9]. Although we recovered virus from genomic constructs containing HAdV-64 HVR1 and HVR7 in the HAdV-5 background, we failed to recover the reverse chimeras. Placing HAdV-64 HVR1 into HAdV-5 increased the HD5 IC_50_ of the virus ∼5-fold over HAdV-5, which was also significantly higher than the HD5 IC_50_ of the pool of viruses from round 50 of the selection (Fig. 2B). In contrast, placing the HAdV-64 HVR7 into HAdV-5 had no effect. And, replacing both HVR1 and HVR7 in HAdV-5 resulted in an intermediate phenotype compared to the single HVR changes. Overall, these studies identify HVR1 as a novel determinant for HAdV-5 neutralization by HD5.

### Multiple sequence elements in penton base contribute to HD5 sensitivity

The absence of mutations in fiber and PB in the evolved viral populations was unexpected based on our previous studies [9]. To substantiate our prior findings and more narrowly delineate neutralization determinants in PB, we created a series of HAdV-5-based chimeric viruses in which portions of PB were swapped with the corresponding residues from HAdV-64. These constructs also contain a DTET to GYAR mutation in fiber, which acts in concert with changes in PB to ablate HD5-dependent neutralization [9]. Initially, we chose highly conserved regions spaced evenly within the HAdV-5 PB sequence as junction points for our chimera designs. As in our prior studies [9], HAdV-5 infection was potently neutralized, HAdV-64 infection was enhanced 4- to 5-fold, and infection by a chimera (C1) containing the entire HAdV-64 PB was moderately enhanced 2- to 3-fold when incubated with 5 µM or 10 µM HD5 (Fig. 3A). If C-terminal residues 288 to 571 of PB are from HAdV-5 (C5), then the virus is neutralized by HD5. If they are from HAdV-64 (C4), then infection is not neutralized but enhanced. An intermediate phenotype of partial HD5 sensitivity was observed when residues 288 to 434 were derived from HAdV-64 and residues 435 to 571 were derived from HAdV-5 (C2), while the inverse construct (C3) completely recapitulated the HD5-sensitivity of HAdV-5. Thus, residues 288 to 434 from HAdV-5 are necessary for complete neutralization, while residues 435-571 also contribute.

**Figure 3.**
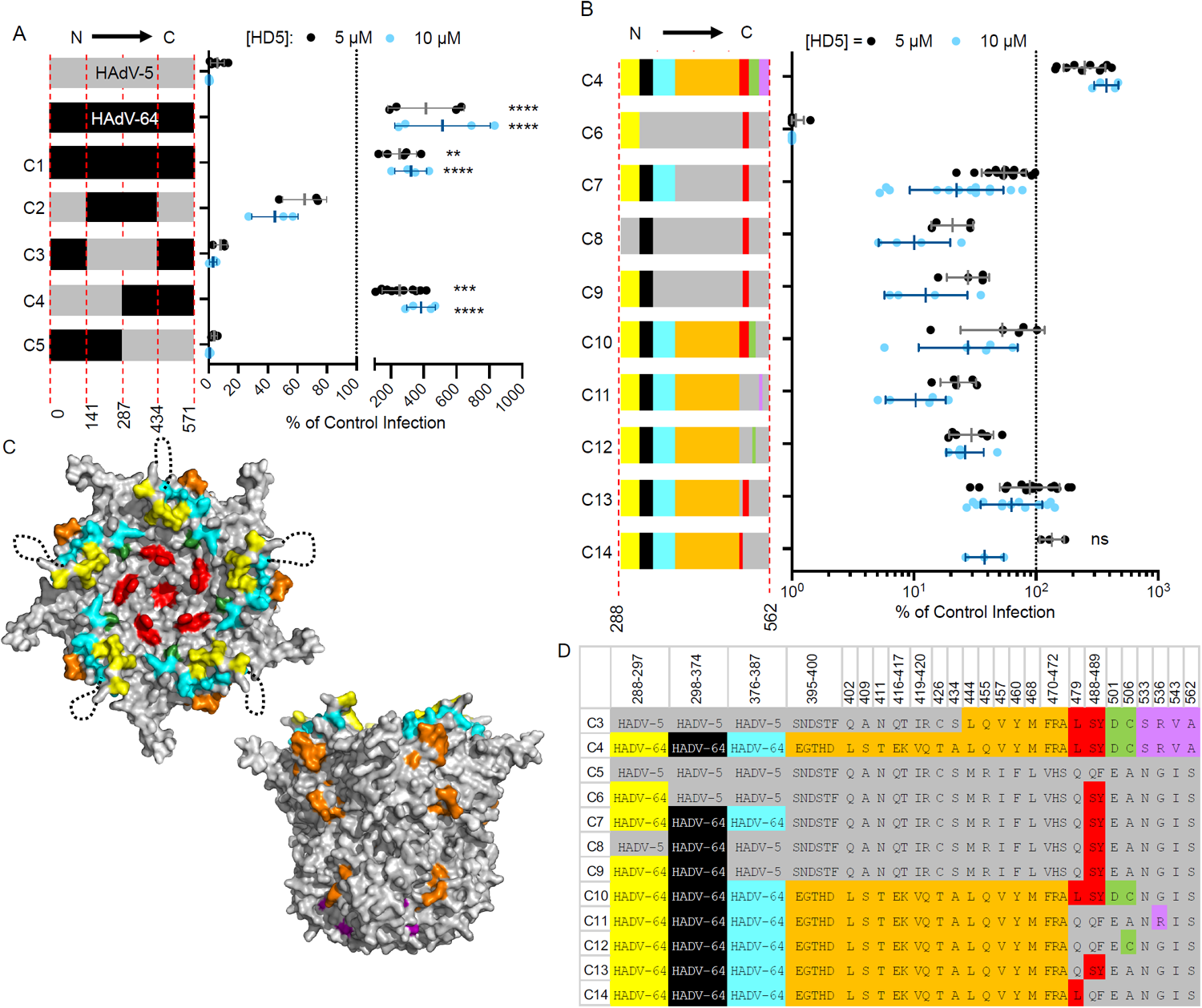
Determinants of HD5-mediated neutralization are located within the C-terminal half of penton base. (A and B) HAdV-5, HAdV-64, and chimeric viruses were incubated with 5 µM (black) or 10 µM (blue) HD5 and then assessed for infectivity on A549 cells. In (A), colors denote sequences derived from HAdV-5 (grey) or HAdV-64 (black), and amino acid residue numbers refer to HAdV-5 PB. In (B), colors correspond to the chart in (D) of the differential amino acid residues between HAdV-5 and HAdV-64 in the C-terminal half of PB, numbered according to HAdV-5. Each data point is an independent experiment, and bars are the mean ± SD of the percent infectivity compared to control cells infected with each virus in the absence of HD5. Note that the same data for C4 is plotted on both graphs. For (A), results of two-way ANOVA with Dunnett’s multiple comparison to HAdV-5 are indicated by asterisks. For (B), all comparisons to chimera C4 are significant (P ≤ 0.01) except where indicated as not significant (ns). (C) Space-filling model of the crystal structure of a pentamer of HAdV-5 PB (PDBID:6CGV [36]) in top and side views, with differential residues colored as in (D). The unresolved RGD loop (residues 298-374) is denoted by a dotted black line.

Although not exhaustive, additional chimeras were created to probe the contribution of specific variable sequences within the C-terminal half of PB to HD5-dependent neutralization (Figs. 3B and 3D). In the description that follows and in Fig 3B-D, colors indicate where the equivalent residues from HAdV-64 are substituted into HAdV-5 PB. The highly variable RGD loop (black) from HAdV-5 is necessary for HD5 neutralization. A moderately conserved motif C-terminal to the RGD loop (cyan) also contributes to HD5 sensitivity, while one N-terminal to the RGD loop (yellow) does not (compare C6, C7, C8, and C9). And, the four variable positions between residues 533 and 562 (purple) contribute to neutralization (compare C4 and C10), while the 22 variable residues between 395 and 472 (orange) do not (compare C7 and C13). Although no definitive conclusions can be drawn, a comparison of C10 and C11 suggests that the variable positions between residues 479 and 506 (red and green) also contribute modestly to neutralization. Collectively, this analysis suggests that complete neutralization of HAdV-5 by HD5 reflects the additive effects of multiple residues in PB.

### The role of capsid proteins in determining the outcome of HD5-virus interactions depends on the timing of exposure to HD5 relative to cell binding

Rational design and directed evolution implicated different major capsid proteins as HD5 neutralization determinants. We suspected that this discrepancy was protocol-dependent. We therefore determined the HD5 IC_50_ for key viruses by either pre-incubating virus with HD5 and then adding this mixture to cells, as in the assessment of PB chimeric viruses (protocol 1), or by adding the defensin to virus pre-bound to cells, as in the selection for HD5-resistance (protocol 2). The phenotypes of HAdV-5 (Fig. 4A), the round 50 pool (Fig. 4B), and hexon HVR1 chimera (Fig. 4C) were largely protocol-independent. The only difference in the phenotype of HAdV-64 (Fig. 4D) was enhancement in protocol 1 compared to resistance in protocol 2. In contrast, the phenotype of the vertex C4 chimera was dramatically protocol-dependent (Fig 4E), demonstrating that cell binding can alter HD5 sensitivity.

**Figure 4.**
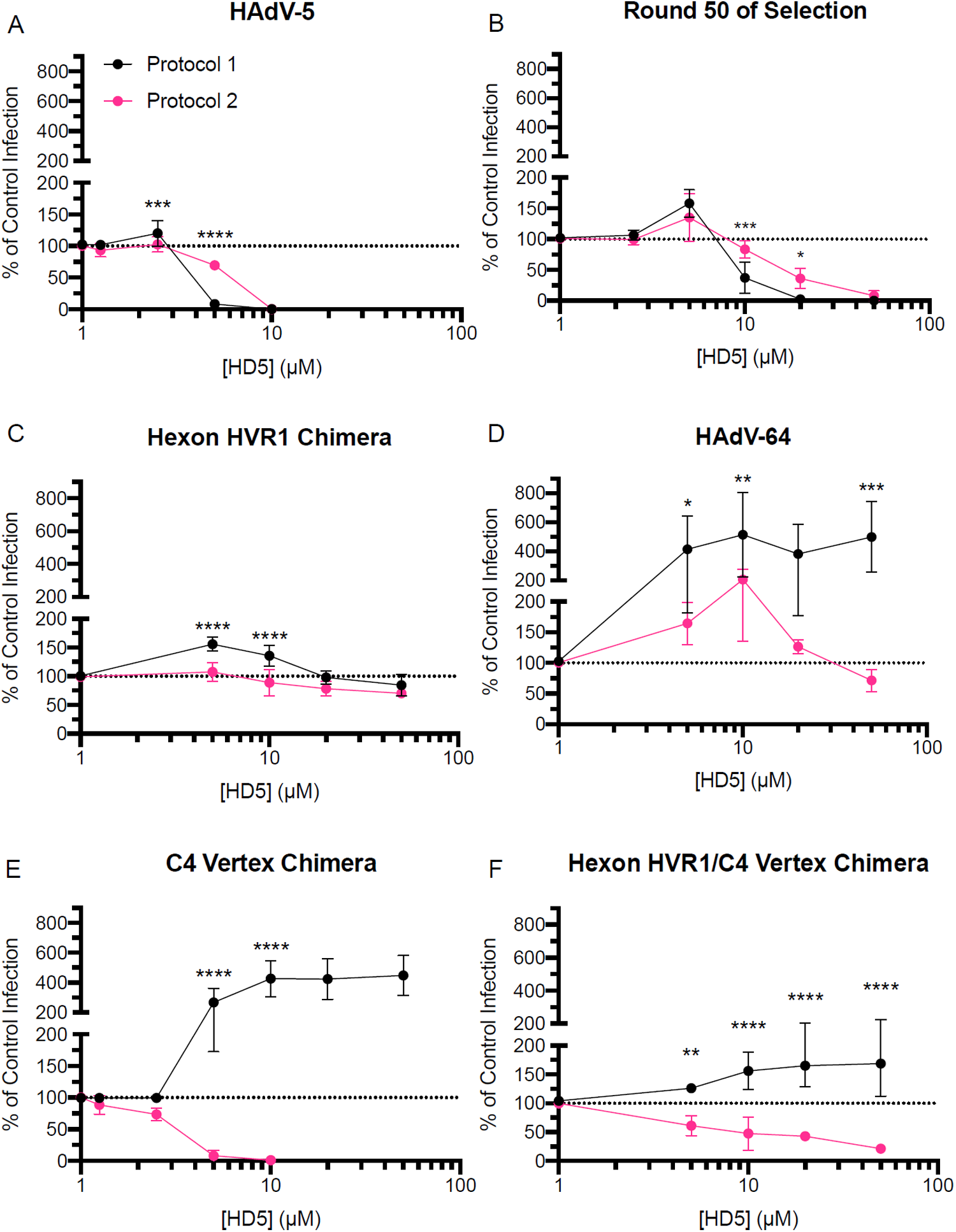
Hexon and vertex play different roles during virus-defensin interactions. Purified (A) HAdV-5, (B) virus expanded from round 50 of selection, (C) hexon HVR1 chimera, (D) HAdV-64, (E) C4 vertex chimera and (F) hexon HVR1/C4 vertex chimera were either incubated with HD5 and then added to A549 cells (protocol 1 – black) or bound to A549 cells prior to HD5 addition (protocol 2 – pink). Data is the mean of 3 to 11 independent experiments ± SD. Results of two-way ANOVA with Sidak’s multiple comparisons at each HD5 concentration are indicated by asterisks.

We then created and tested an additional construct combining the changes from the C4 vertex and hexon HVR1 chimeras. In protocol 1, the hexon HVR1/C4 vertex chimera had the same phenotype as the hexon HVR1 chimera rather than the C4 vertex chimera: it was resistant to neutralization by HD5 at all concentrations but not enhanced (Fig 4F). In protocol 2, the hexon HVR1/C4 vertex chimera exhibited an intermediate phenotype: it was ∼3-fold more HD5-resistant than the C4 vertex chimera but ∼2-fold more HD5-sensitive than the hexon HVR1 chimera. Thus, both hexon HVR1 and the vertex are important determinants of viral infectivity in the presence of HD5, but they do not act synergistically. Rather, the phenotype of HVR1 is predominant in protocol 1, while HVR1 and the vertex appear to have additive effects in protocol 2.

### Vertex and hexon proteins dictate HD5 binding to the viral capsid

Based on previous studies demonstrating a direct interaction between HD5 and HAdV [9, 11, 12, 15], we quantified the average number of HD5 molecules bound to the capsid of each virus. At a concentration of 20 µM, HD5 binds at a high molar ratio to HAdV-5 (7090 +/- 1550 molecules of HD5 per HAdV-5 virion), as shown previously [9], and ∼83-fold less to HAdV-64 (Fig 5A). Interestingly, the C4 vertex chimera bound ∼2.3-fold more than HAdV-5, while the hexon HVR1 chimera bound ∼5-fold less. Finally, HD5 bound to the combined hexon HVR1/C4 vertex chimera at the same levels as HAdV-5. At lower concentrations (5 µM and 10 µM), HD5 binding to HAdV-5, C4 vertex chimera, and hexon HVR1/C4 vertex chimera was equivalent (Fig 5B). Thus, changes in both hexon and the vertex proteins alter the stoichiometry of the HD5-capsid interaction.

**Figure 5.**
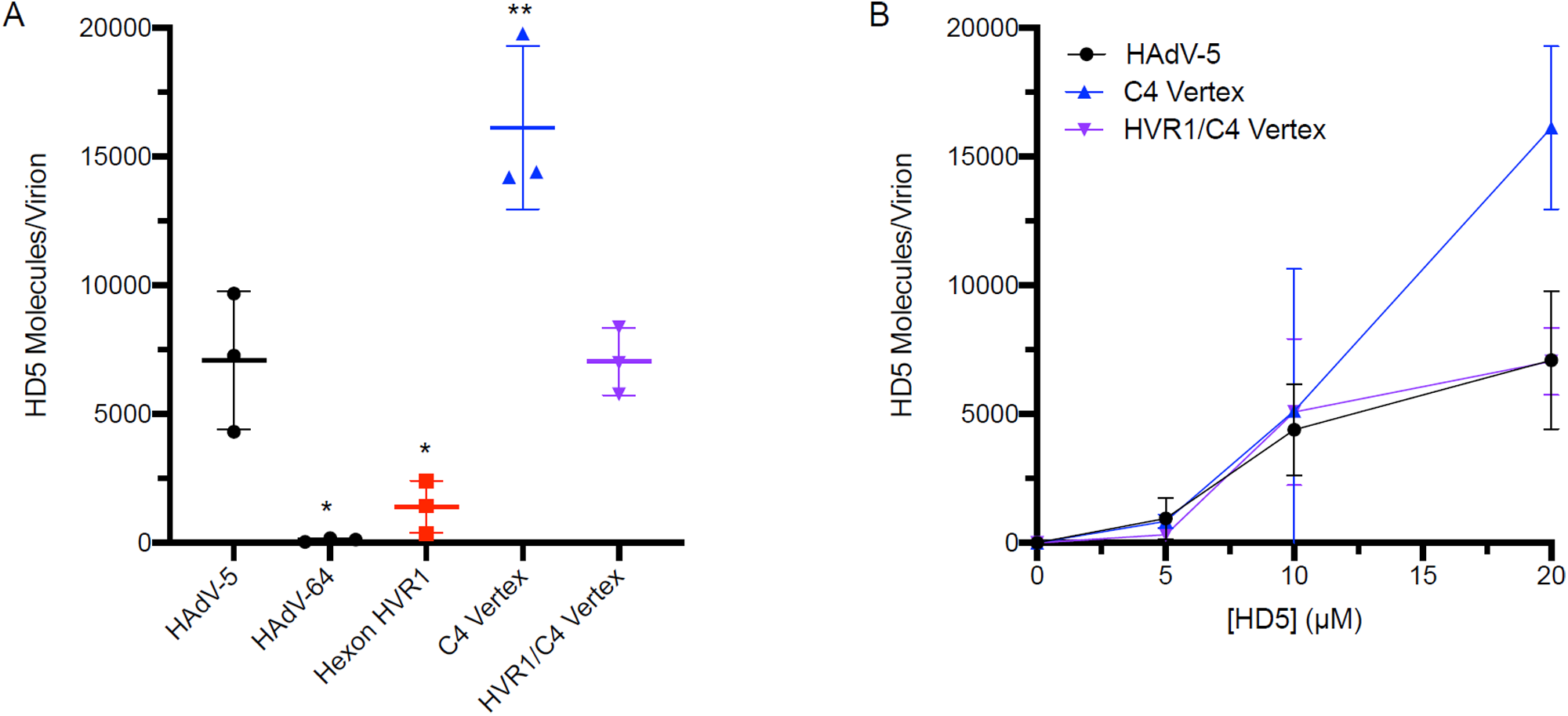
Both hexon HVR1 and vertex contribute to HD5 binding. (A) HAdV-5, HAdV-64, and chimeric viruses were incubated in the presence of 20 µM HD5. The amount of HD5 bound per virion was quantified from 3 independent experiments. (B) HD5 molecules bound per virion of HAdV-5 and chimeras C4 vertex and hexon HVR1/C4 vertex were also determined in the presence of 5 µM and 10 µM HD5. Note that the data for 20 µM HD5 for these viruses in (A) is reproduced in (B). Lines are the mean ± SD. The results of ordinary one-way ANOVA with Dunnett’s multiple comparisons to HAdV-5 is denoted by asterisks.

### The composition of both hexon HVR1 and the vertex proteins influence fiber stability upon HD5 binding

Our prior studies are consistent with a mechanism in which HD5 neutralizes HAdV-5 by stabilizing the capsid and preventing shedding of the vertex proteins [9, 11]. Thus, capsid changes could impact the thermostability of the virus in the presence or absence of HD5. We first determined the temperature at which 50% of fiber dissociates from the viral capsid (Tm) in the absence of HD5. As expected, all viruses remained intact at 44°C, and fiber was completely dissociated at 50°C (Fig 6A and B). The Tm of the hexon HVR1 chimera was identical to that of HAdV-5 (46°C), while the Tms of the C4 vertex and hexon HVR1/C4 vertex chimeras were similar to each other and warmer than that of HAdV-5 by ∼2°C. Thus, amino acid changes in the C4 vertex stabilized the fiber-capsid interaction, while amino acid changes in hexon HVR1 had no effect. We then tested the effect of HD5 on fiber dissociation, when samples were heated to 2°C above the Tm to assure full fiber dissociation in the absence of HD5. Despite their different infection phenotypes and HD5 binding capacities, the fiber of each of these viruses was fully capsid-associated upon incubation with 20 µM HD5 (Fig 6C). HAdV-5 and C4 vertex chimera fibers were 50% capsid-associated at 5 µM HD5 and had identical HD5-dependent dissociation profiles. The hexon HVR1/C4 vertex chimera required a 2-fold lower HD5 concentration than HAdV-5 to be 50% capsid-associated and was significantly more stabilized than HAdV-5 at both 2.5 µM and 5 µM HD5. In contrast, the hexon HVR1 chimera required at least a 2-fold higher HD5 concentration than HAdV-5 to be 50% stabilized. Overall, the composition of both HVR1 and the vertex influence HD5-mediated fiber stabilization; however, the phenotype of the combination does not mirror the individual contributions of each capsid component.

**Figure 6.**
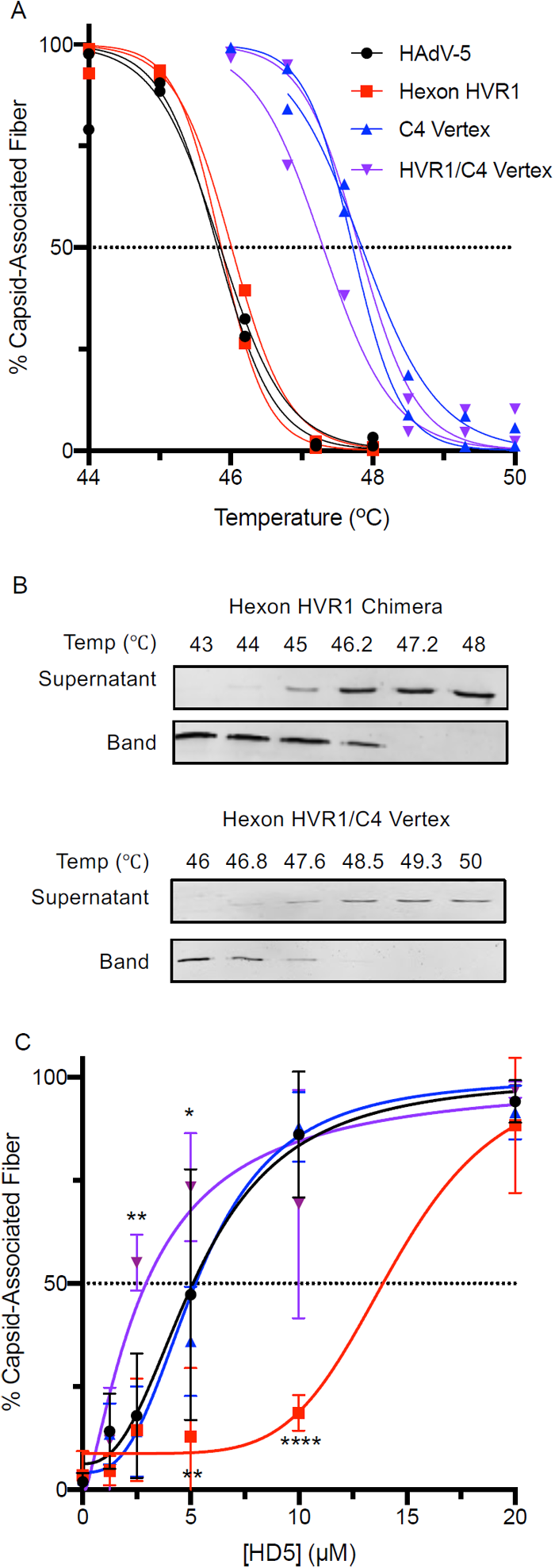
Fiber thermostability does not correlate with infection phenotype. The percent of fiber that remains capsid associated was determined (A) as a function of temperature in the absence of HD5 or (C) as a function of HD5 concentration for HAdV-5 (black) and hexon HVR1 chimera (red) at 48°C and for C4 vertex chimera (blue) and hexon HVR1/C4 vertex chimera (purple) at 49.3°C. (B) Representative immunoblots from the temperature gradients of hexon HVR1 and hexon HVR1/C4 vertex chimeras are shown. In (A) each point and line is an individual replicate. In (C), each point is the mean ± SD of 3 independent experiments, and the results of two-way ANOVA with Dunnett’s multiple comparisons to HAdV-5 is denoted by asterisks.

## Discussion

In this study, we have directly demonstrated that enteric α-defensins can impose selective pressure on non-enveloped viral evolution. We postulated that selection may occur during fecal-oral transmission due to the abundant expression and high concentration of enteric α-defensins in the intestinal lumen [16]. This hypothesis is based on the observation that naturally occurring human and mouse AdVs are differentially susceptible to α-defensin antiviral activity and that defensin resistance correlates with AdV species that contain fecal-orally transmitted serotypes [9, 16, 18, 28]. It would also explain the α-defensin resistance of echovirus and reovirus [17]. A prior study used a similar approach to select isolates of HIV-1, an enveloped virus, resistant to retrocyclin, a θ-defensin expressed in non-human primates [29]. The mutations that appeared in both HAdV-5 and HIV-1 during selection increased the positive charge of the viral structural proteins (hexon and gp41, respectively), likely leading to an overall decrease in defensin binding due to the cationicity of defensins. A similar principle of surface charge modulation contributes to the experimental evolution of bacterial resistance to cationic antimicrobial peptides, which also confers resistance to β-defensins [30]. Collectively, there is now direct evidence that defensins can impose selective pressure on the evolution of a wide range of organisms including not only bacteria and enveloped viruses but non-enveloped viruses as well.

Another major outcome of our investigation is the identification of a novel determinant of HAdV-5 neutralization by HD5 that fundamentally alters our understanding of the α-defensin antiviral mechanism. Our prior studies supported a model where HD5 binds to the fiber and PB proteins at the vertices of the HAdV-5 capsid and stabilizes their interaction [9]. This action blocks uncoating of the capsid during cell entry and restricts release of the membrane-lytic protein VI, which in turn prevents endosome escape and precludes trafficking of the viral genome to the nucleus [11, 14]. This mechanism was supported by structural, biochemical, biophysical, and genetic studies [9, 11, 12, 15, 31]; however, we were unable to account for the extensive HD5 binding to the hexons of HAdV-5 in our cryoEM studies [9]. Directed evolution of HAdV-5 under selection by HD5 has provided new insight into the role of HD5-hexon interactions that substantially revises this model.

A newly identified role for hexon is to mediate HD5-enhanced infection. We define enhancement as ≥2-fold higher infection in the presence of HD5 than in the absence of HD5. Consistent with prior studies demonstrating that defensin interactions with the HAdV capsid promote cell binding [9, 15], we found that enhancement only occurs when the virus binds HD5 before binding to the cell (protocol 1 in Figs. 3 and 4). Increased cell binding likely occurs by neutralizing the repulsive forces of the electronegative capsid in proximity to the cell membrane [15], although HD5 could also bridge interactions between the virus and cellular lipids, glycans, or an unidentified HD5-specific receptor. A similar mechanism has been shown in shigella, another gastrointestinal pathogen that appropriates HD5 to facilitate infection [19, 20]. In addition, only viruses containing hexons in which all of the HVRs are from one serotype are enhanced, including a recently published HAdV-5 chimera containing all of the HVRs of HAdV-48 hexon [32]. This suggests that homotypic interactions involving hexon HVRs mediate enhancement. A comparable role for hexon in mediating cell binding has been previously described, where coagulation factor X (FX) bridges an interaction between the virus and heparan sulfate proteoglycans on hepatocytes [33, 34]; however, HD5 and FX target distinct hexon HVRs. Thus, our studies have identified a novel role for hexon as the sole determinant of cell binding leading to defensin-enhanced infection.

The HVR1 loop of hexon also functions in cooperation with the vertex proteins as a previously unidentified determinant of HD5-mediated neutralization. Despite increased cell binding and the potential for enhanced infection, many HAdV serotypes are nonetheless neutralized by HD5 [9]. If either HVR1 or both vertex determinants (four residues near the N-terminus of fiber and the C-terminal half of PB) are derived from a resistant virus, then neutralization does not occur. This suggests that the functions of the capsid determinants are interrelated. However, if the virus is bound to its cellular receptor and co-receptor prior to HD5 addition (protocol 2 in Figs. 1C and 4), HVR1 is the sole determinant of neutralization. Thus, the virus-cell interaction functionally replaces the vertex in potentiating HD5 neutralization. Mechanistically, this could occur through receptor-induced conformational changes that lead to exposure of HD5-interacting surfaces in the vertex that are buried in the absence of receptor. Alternatively, the receptor-virus interface could provide a novel target for HD5 binding. However, these interpretations suggest that the virus in protocol 1 either doesn’t experience the conformation induced at 4°C in protocol 2 or transitions through it too rapidly for HD5 to exert a neutralizing effect. Moreover, it is unknown how many vertices are receptor-engaged under the conditions of protocol 2. If only a subset are bound, then blocking uncoating triggered through these vertices may be the key step impeded by HD5 binding [35]. The nature of the HVR1 loop also dictates the Hill slope of the HD5 inhibition curve in protocol 2 (Fig 4), suggesting distinct levels of cooperativity and modes of HD5 binding by capsids differing in HVR1 loops. And, C4 vertex-containing viruses are neutralized at a lower HD5 concentration than those with the HAdV-5 vertex, which may be due to the inherently higher thermostability of the C4 vertex (Fig. 6A) or to its higher HD5-binding capacity (Fig. 5). Although a minimum amount of HD5 binding to the capsid is required for neutralization, there is not a simple correlation between the degree of neutralization and the amount of HD5 bound. Total HD5 bound appears to reflect additive functions of HVR1 and the vertex. And, neither the HAdV-5 nor HAdV-64 vertex appears to bind HD5 to the same extent as the C4 vertex, suggesting altered binding by the artificial interface in the C4 chimera. We also found that fiber stabilization does not directly correlate with the infection phenotypes of the chimeras and that swapping HVR1 also affected the ability of HD5 to stabilize fiber dissociation. Collectively, these findings suggest that our previous model of vertex stabilization mediated only by HD5 interactions with fiber and PB is incomplete.

A model most consistent with our data is that blocking vertex dissociation through HD5 interactions with fiber/PB is insufficient, and a separate hexon-dependent mechanism, perhaps inter- or intra-hexon or hexon-vertex “cross-linking” by HVR1-HD5 interactions, is also required to prevent uncoating. We cannot formally exclude a model where the capsid determinants act cooperatively to coordinate HD5 binding at the vertex, particularly since the HAdV-5 hexon HVR1 loop is long enough to extend from the peri-pentonal hexons towards fiber. However, in that model it is harder to rationalize a role for the point mutation that arose in hexon HVR7 in the later rounds of selection. Both models are also consistent with the phenotypes of the PB chimeras, where intermediate levels of neutralization result from a subset of the PB changes found in the C4 vertex. Further experimentation will be required to resolve these possibilities.

In summary, our studies generated two major insights. First, we proved that HD5 can act as a selective pressure on the evolution of non-enveloped viruses. Second, we identified hexon as a key contributor to α-defensin interactions with AdV. Importantly, all three HAdV capsid proteins play a role in neutralization, but hexon rather than PB or fiber determines enhancement. Thus, we have extensively revised the prior model of HD5-mediated neutralization and established the feasibility of this process to shape viral evolution *in vivo*, which is further supported by the recent demonstration of neutralization and enhancement of HAdV-based vectors by HD5 in a mouse model [32].

## Materials and Methods

### Cell lines

HEK 293 cells overexpressing human β5 integrin (293β5) [9] and A549 cells (ATCC) were maintained in Dulbecco’s modified Eagle’s medium (DMEM) with 10% fetal bovine serum (FBS), penicillin, streptomycin, _L_-glutamine, and non-essential amino acids (complete media).

### HD5

Partially purified (89%) linear peptides (UniProtKB: ATCYCRTGRCATRESLSGVCEISGRLYRLCCR) were synthesized (LifeTein, Somerset, NJ), oxidatively folded, and purified by reverse-phase high-pressure liquid chromatography (RP-HPLC) [12]. Fractions containing the correctly folded species were lyophilized, resuspended in deionized water, and quantified by absorbance at 280 nm as described [12]. Purity (>99%) and mass were verified by analytical RP-HPLC and MALDI-TOF mass spectrometry. HD5 was stored at −80°C.

### Viruses

An E1/E3-deleted, replication-defective HAdV-5 vector containing a CMV promoter-driven enhanced green fluorescent protein (eGFP) reporter gene cassette was used as the parent construct for all of the novel chimeras created for these studies, which were generated by recombineering in BACmids [9, 38]. The C1 chimera was previously referred to as “PB/GYAR” [9]. Designs of chimeras C2-C14 are depicted in Fig. 3. Hexon chimeras were created by replacing either HVR1 (bp 19247 to 19336 in HAdV-5, NCBI: AC_000008.1), HVR7 (bp 20090 to 20203), or both HVR1 and HVR7 in the HAdV-5 *hexon* ORF with the corresponding sequences from HAdV-64 HVR1 (bp 18196 to 18243, GenBank: EF121005.1) and HVR7 (bp 19027 to 19161). The combined hexon HVR1/C4 vertex chimera was created by replacing the HAdV-5 HVR1 with that of HAdV-64 in the C4 chimera. A previously described E3 deleted, replication-competent HAdV-64 virus containing a CMV-eGFP ORF was also used to study the effects of HD5 on infection of a virus with a WT HAdV-64 capsid [39]. The fidelity of all BACmid constructs was verified by Sanger sequencing of the recombineered region and by restriction digest of the entire BACmid. In addition, the BACmids of HAdV-64, chimera C1, and chimera C4 were sequenced in their entirety by whole genome sequencing.

To produce virus, 293β5 cells were transfected with viral genomes released from the BACmids by *Pac* I endonuclease digestion. Following amplification over several passages in 293β5 cells to generate sufficient inoculum, approximately eight to ten T175 flasks of 293β5 cells were infected at a multiplicity of infection of ∼3. Upon development of complete cytopathic effect, virus was precipitated from the supernatant in 8% polyethylene glycol (PEG) [40]. Virus was then purified from cell lysates and the PEG precipitate using a CsCl gradient as previously described [14]. Purified virus was dialyzed against three changes of 150 mM NaCl, 40 mM Tris, 10% glycerol, 2 mM MgCl_2_, pH 8.1, snap frozen in liquid nitrogen, and stored at −80°C. The viral particle concentration was determined by Qubit fluorometric quantification (ThermoFisher) against a DNA standard (1 µg = 2.34E+10 virions). Genomic DNA was isolated from purified virus using the GeneJET genomic DNA purification kit (Thermo Fisher), and the fidelity of the changed regions was verified by Sanger sequencing. For biochemical assays, viral protein concentration was determined by Bio-Rad Protein Assay with a bovine serum albumin standard.

### Infection Assays

As in our previous studies [15], we employed two protocols, which differed in the order of addition of HD5 to the virus relative to cell binding. All infection assays were performed in black wall, clear bottom 96-well plates seeded with either A549 or 293β5 cells. Protocol 1 (Figs. 2, 3, and 4): virus and HD5 were incubated together on ice in serum free media (SFM) for 45 min. Cells were then washed twice with SFM to remove any residual serum, the virus/HD5 mixture was added to the cells, and the plate was shifted to 37°C. Protocol 2 (Figs. 1C, and 4): cells were incubated on ice for 5 min, washed once with cold SFM, and virus in SFM was then added. After incubation for 45 min on ice, the inoculum was removed, the cells were washed once with SFM, and HD5 in SFM was added. After incubation for 45 min on ice, the plate was shifted to 37°C. For both methods, the cells were washed with SFM after 2 h of incubation at 37°C, and the media was replaced with complete media made from phenol red free DMEM. Cells were imaged 20-28 h post-infection on a Typhoon (GE Healthcare) or Sapphire (Azure) imager, and ImageJ was used to quantify background-subtracted total monolayer fluorescence. Data are shown as a percent of control infection in the absence of HD5. Concentrations of each virus were determined in advance that result in 50-80% of maximum signal for inhibition studies or 10-25% of maximum signal for enhancement studies.

### HD5 Selection

As previously described, we used purified “mutator” adenovirus (HAdV-5.polF421Y) that was passaged for 10 rounds in 293β5 cells to generate diversity in the starting population [24]. We infected 293β5 cells in 12-well plates under selection with HD5 at the IC_90_ following protocol 2 above. We used a ∼6-fold lower MOI for control virus passaged in the absence of HD5 to yield comparable infection levels. Upon development of complete cytopathic effect, which typically occurred 3 to 4 days post-infection, the entire culture of cells and media was collected. A clarified freeze/thaw lysate generated from the infected cells was combined with the supernatant and snap frozen in liquid nitrogen for storage at −80°C. An aliquot was used to determine the infectious titer of each sample to ensure a similar level of infection for each round of selection. In addition, the HD5 IC_90_ was determined periodically on 293β5 cells to recalibrate the selective pressure (Fig. 1A). For subsequent assays, selected viral pools were amplified in the absence of HD5 over ∼5 passages in 293β5 cells to generate sufficient inoculum and then purified from preparations of eight to ten T175 flasks of 293β5 cells as described above.

### Plaque Purification

Pooled virus from rounds 10, 40, and 50 of the selection were plaque purified on 293β5 cells in a 6-well plate. For the initial infection, purified virus was used to infect cells at low MOI (0.004 - 0.03), and cultures were overlaid with complete media containing 1% Difco Noble Agar. After 7-14 d, cells and agar plugs from individual plaques were harvested using a pipet tip, resuspended in 100 µL of complete media, and lysed through three freeze-thaw cycles. The plaque-purified isolates were subjected to two additional rounds of plaque purification, expanded, and purified as described above. At intermediate steps, PCR and Sanger sequencing were used to identify plaques containing mutations in the *hexon* and *L4-100K* ORFs.

### Whole Genome Sequencing

Genomic DNA was extracted and quantified from purified preparations of the plaque purified viruses and the pooled viruses from every 5^th^ round of HD5 selection and the 10^th^ and 20^th^ round of control selection. The Nextera XT DNA Library preparation kit (Illumina) was used to tagment and barcode the genomic DNA. A MiSeq v3 150 cycle reagent kit (Illumina) was used to sequence the libraries with 75 base paired-end reads. The data was analyzed using the BreSeq pipeline to align the sequences to the parent genome and identify mutations [25].

### Virus-Defensin Binding Assay

To measure HD5 binding, 2.5 µg of purified virus was incubated with 5, 10, or 20 µM HD5 in a buffer consisting of 150 mM NaCl, 20 mM Tris, 5% glycerol, 1 mM MgCl_2_ on ice for 45 min. Samples were then layered onto a discontinuous gradient containing 300 µl of 30% Histodenz overlaying 200 µl of 80% Histodenz in 20 mM tris pH 7.4. Gradients were centrifuged using an SW55ti rotor with adaptors (Beckman) at 209,0006 × g (avg.) for 1.5 h at 4°C, and the visible virus band was collected. Samples containing HAdV-5 mixed with HD5 that were not subject to centrifugation were used to generate a standard curve for quantification. All samples were reduced with DTT, heated to 95°C for 5 min, and separated by SDS-PAGE (10-20% tris-tricine gel). The gels were stained with Flamingo fluorescent protein gel stain (Bio-Rad) and imaged on a Sapphire Biomolecular Imager (Azure). Protein bands were quantified using Azure Spot software (Azure). The amount of HD5 in each sample was normalized to protein V and protein VII and quantified against the standard curve using Prism 8.3.0 software (GraphPad).

### Thermostability Assay

To measure the capsid association of fiber, 250 ng of purified virus was incubated on ice with or without the indicated concentrations of HD5 for 45 min in serum free DMEM containing 0.05% BSA, 150 mM NaCl, and 10 mM HEPES pH 7.5. Samples were heated in a thermocycler at the indicated temperatures for 10 min and loaded onto a discontinuous gradient containing 400 µl of 30% Histodenz and 200 µl of 80% Histodenz in 20 mM tris pH 7.4. Samples were centrifuged as described above. Fractions were collected as follows: 90 µl from the top (supernatant), 2 middle fractions of 150 µl, and then 90 µl (virus band). The supernatant and band fractions were reduced with DTT, heated to 95°C for 5 min, separated by SDS-PAGE (12% tris-glycine gel), transferred to nitrocellulose, and probed by immunoblot for fiber using the 4D2 monoclonal antibody (ThermoFisher) and an Alexa Fluor 647-conjugated secondary antibody. Blots were imaged and quantified as described above to determine the fraction of total fiber in each sample that was present in the virus band fraction. Prism 8.3.0 was used for non-linear regression analysis to calculate Tm and HD5 concentrations resulting in 50% fiber dissociation in Fig. 6.

### Statistical analysis and structural rendering

Statistical analysis was performed using Prism 8.3.0. Specific analyses are indicated in the figure legends. For all tests, not significant (ns), P > 0.05; *, P = 0.01 to 0.05; **, P = 0.001 to 0.01; ***, P = 0.0001 to 0.001; ****, P < 0.0001. Structural figures were generated using the PyMOL Molecular Graphics System, Version 2.0.7 Schrödinger, LLC.

## Data availability statement

The data that support the findings of this study are available from the corresponding author upon reasonable request.

## Acknowledgements

We thank Carissa M. Lucero for assistance with the selection. We also thank Rossana Colón-Thillet for assistance with recombineering the virus used in Fig. 2A column 10. Research reported in this publication was supported by the National Institute of Allergy and Infectious Diseases of the National Institutes of Health (NIH) under awards K22AI081870, R01AI104920, T32AI083203, and F30AI140620 and by the Office of the Director of the NIH under award S10OD026741. The funders had no role in the study design, data collection and analysis, decision to publish, or preparation of the manuscript.

## Notes

### Competing Interest Statement

The authors have declared no competing interest.

## References

1. Holly MK, Diaz K, Smith JG. Defensins in Viral Infection and Pathogenesis. Annu Rev Virol. 2017;4(1):369-91. Epub 2017/07/19. doi: 10.1146/annurev-virology-101416-041734. PubMed PMID: 28715972.

2. Selsted ME, Ouellette AJ. Mammalian defensins in the antimicrobial immune response. Nat Immunol. 2005;6(6):551-7. Epub 2005/05/24. doi: 10.1038/ni1206. PubMed PMID: 15908936.

3. Wilson SS, Wiens ME, Smith JG. Antiviral mechanisms of human defensins. J Mol Biol. 2013;425(24):4965-80. Epub 2013/10/08. doi: 10.1016/j.jmb.2013.09.038. PubMed PMID: 24095897; PubMed Central PMCID: PMCPMC3842434.

4. Brice DC, Diamond G. Antiviral Activities of Human Host Defense Peptides. Curr Med Chem. 2019. Epub 2019/08/07. doi: 10.2174/0929867326666190805151654. PubMed PMID: 31385762.

5. Wiens ME, Smith JG. Alpha-defensin HD5 inhibits furin cleavage of human papillomavirus 16 L2 to block infection. J Virol. 2015;89(5):2866-74. Epub 2014/12/30. doi: 10.1128/JVI.02901-14. PubMed PMID: 25540379; PubMed Central PMCID: PMCPMC4325740.

6. Wiens ME, Smith JG. alpha-Defensin HD5 Inhibits Human Papillomavirus 16 Infection via Capsid Stabilization and Redirection to the Lysosome. mBio. 2017;8(1). Epub 2017/01/26. doi: 10.1128/mBio.02304-16. PubMed PMID: 28119475; PubMed Central PMCID: PMCPMC5263252.

7. Dugan AS, Maginnis MS, Jordan JA, Gasparovic ML, Manley K, Page R, et al. Human alpha-defensins inhibit BK virus infection by aggregating virions and blocking binding to host cells. J Biol Chem. 2008;283(45):31125-32. Epub 2008/09/11. doi: 10.1074/jbc.M805902200. PubMed PMID: 18782756; PubMed Central PMCID: PMCPMC2576552.

8. Zins SR, Nelson CD, Maginnis MS, Banerjee R, O’Hara BA, Atwood WJ. The human alpha defensin HD5 neutralizes JC polyomavirus infection by reducing endoplasmic reticulum traffic and stabilizing the viral capsid. J Virol. 2014;88(2):948-60. Epub 2013/11/08. doi: 10.1128/JVI.02766-13. PubMed PMID: 24198413; PubMed Central PMCID: PMCPMC3911681.

9. Smith JG, Silvestry M, Lindert S, Lu W, Nemerow GR, Stewart PL. Insight into the mechanisms of adenovirus capsid disassembly from studies of defensin neutralization. PLoS Pathog. 2010;6(6):e1000959. Epub 2010/06/30. doi: 10.1371/journal.ppat.1000959. PubMed PMID: 20585634; PubMed Central PMCID: PMCPMC2891831.

10. Buck CB, Day PM, Thompson CD, Lubkowski J, Lu W, Lowy DR, et al. Human alpha-defensins block papillomavirus infection. Proc Natl Acad Sci U S A. 2006;103(5):1516-21. Epub 2006/01/25. doi: 10.1073/pnas.0508033103. PubMed PMID: 16432216; PubMed Central PMCID: PMCPMC1360544.

11. Smith JG, Nemerow GR. Mechanism of adenovirus neutralization by Human alpha-defensins. Cell Host Microbe. 2008;3(1):11-9. Epub 2008/01/15. doi: 10.1016/j.chom.2007.12.001. PubMed PMID: 18191790.

12. Tenge VR, Gounder AP, Wiens ME, Lu W, Smith JG. Delineation of interfaces on human alpha-defensins critical for human adenovirus and human papillomavirus inhibition. PLoS Pathog. 2014;10(9):e1004360. Epub 2014/09/05. doi: 10.1371/journal.ppat.1004360. PubMed PMID: 25188351; PubMed Central PMCID: PMCPMC4154873.

13. Gulati NM, Miyagi M, Wiens ME, Smith JG, Stewart PL. alpha-Defensin HD5 Stabilizes Human Papillomavirus 16 Capsid/Core Interactions. Pathog Immun. 2019;4(2):196-234. Epub 2019/10/05. doi: 10.20411/pai.v4i2.314. PubMed PMID: 31583330; PubMed Central PMCID: PMCPMC6755940.

14. Nguyen EK, Nemerow GR, Smith JG. Direct evidence from single-cell analysis that human {alpha}-defensins block adenovirus uncoating to neutralize infection. J Virol. 2010;84(8):4041-9. Epub 2010/02/05. doi: 10.1128/JVI.02471-09. PubMed PMID: 20130047; PubMed Central PMCID: PMCPMC2849482.

15. Gounder AP, Wiens ME, Wilson SS, Lu W, Smith JG. Critical determinants of human alpha-defensin 5 activity against non-enveloped viruses. J Biol Chem. 2012;287(29):24554-62. Epub 2012/05/29. doi: 10.1074/jbc.M112.354068. PubMed PMID: 22637473; PubMed Central PMCID: PMCPMC3397880.

16. Wilson SS, Bromme BA, Holly MK, Wiens ME, Gounder AP, Sul Y, et al. Alpha-defensin-dependent enhancement of enteric viral infection. PLoS Pathog. 2017;13(6):e1006446. Epub 2017/06/18. doi: 10.1371/journal.ppat.1006446. PubMed PMID: 28622386; PubMed Central PMCID: PMCPMC5489213.

17. Daher KA, Selsted ME, Lehrer RI. Direct inactivation of viruses by human granulocyte defensins. J Virol. 1986;60(3):1068-74. Epub 1986/12/01. PubMed PMID: 3023659; PubMed Central PMCID: PMCPMC253347.

18. Holly MK, Smith JG. Adenovirus Infection of Human Enteroids Reveals Interferon Sensitivity and Preferential Infection of Goblet Cells. J Virol. 2018;92(9). Epub 2018/02/23. doi: 10.1128/JVI.00250-18. PubMed PMID: 29467318; PubMed Central PMCID: PMCPMC5899204.

19. Xu D, Liao C, Xiao J, Fang K, Zhang W, Yuan W, et al. Human Enteric Defensin 5 Promotes Shigella Infection of Macrophages. Infect Immun. 2019;88(1). Epub 2019/10/16. doi: 10.1128/IAI.00769-19. PubMed PMID: 31611271; PubMed Central PMCID: PMCPMC6921650.

20. Xu D, Liao C, Zhang B, Tolbert WD, He W, Dai Z, et al. Human Enteric alpha-Defensin 5 Promotes Shigella Infection by Enhancing Bacterial Adhesion and Invasion. Immunity. 2018;48(6):1233-44 e6. Epub 2018/06/03. doi: 10.1016/j.immuni.2018.04.014. PubMed PMID: 29858013; PubMed Central PMCID: PMCPMC6051418.

21. Cullen TW, Schofield WB, Barry NA, Putnam EE, Rundell EA, Trent MS, et al. Antimicrobial peptide resistance mediates resilience of prominent gut commensals during inflammation. Science. 2015;347(6218):170–5. doi: 10.1126/science.1260580.

22. Masuda K, Sakai N, Nakamura K, Yoshioka S, Ayabe T. Bactericidal activity of mouse alpha-defensin cryptdin-4 predominantly affects noncommensal bacteria. J Innate Immun. 2011;3(3):315-26. Epub 2010/11/26. doi: 10.1159/000322037. PubMed PMID: 21099205.

23. Salzman NH, Hung K, Haribhai D, Chu H, Karlsson-Sjoberg J, Amir E, et al. Enteric defensins are essential regulators of intestinal microbial ecology. Nat Immunol. 2010;11(1):76-83. Epub 2009/10/27. doi: 10.1038/ni.1825. PubMed PMID: 19855381; PubMed Central PMCID: PMCPMC2795796.

24. Myers ND, Skorohodova KV, Gounder AP, Smith JG. Directed evolution of mutator adenoviruses resistant to antibody neutralization. J Virol. 2013;87(10):6047-50. Epub 2013/03/15. doi: 10.1128/JVI.00473-13. PubMed PMID: 23487468; PubMed Central PMCID: PMCPMC3648140.

25. Deatherage DE, Barrick JE. Identification of mutations in laboratory-evolved microbes from next-generation sequencing data using breseq. Methods Mol Biol. 2014;1151:165-88. Epub 2014/05/20. doi: 10.1007/978-1-4939-0554-6_12. PubMed PMID: 24838886; PubMed Central PMCID: PMCPMC4239701.

26. Cepko CL, Sharp PA. Assembly of adenovirus major capsid protein is mediated by a nonvirion protein. Cell. 1982;31(2 Pt 1):407-15. Epub 1982/12/01. doi: 10.1016/0092-8674(82)90134-9. PubMed PMID: 7159928.

27. Yan J, Dong J, Wu J, Zhu R, Wang Z, Wang B, et al. Interaction between hexon and L4-100K determines virus rescue and growth of hexon-chimeric recombinant Ad5 vectors. Sci Rep. 2016;6:22464. Epub 2016/03/05. doi: 10.1038/srep22464. PubMed PMID: 26934960; PubMed Central PMCID: PMCPMC4776158.

28. Gounder AP, Myers ND, Treuting PM, Bromme BA, Wilson SS, Wiens ME, et al. Defensins Potentiate a Neutralizing Antibody Response to Enteric Viral Infection. PLoS Pathog. 2016;12(3):e1005474. Epub 2016/03/05. doi: 10.1371/journal.ppat.1005474. PubMed PMID: 26933888; PubMed Central PMCID: PMCPMC4774934.

29. Cole AL, Yang OO, Warren AD, Waring AJ, Lehrer RI, Cole AM. HIV-1 adapts to a retrocyclin with cationic amino acid substitutions that reduce fusion efficiency of gp41. J Immunol. 2006;176(11):6900-5. PubMed PMID: 16709850.

30. Andersson DI, Hughes D, Kubicek-Sutherland JZ. Mechanisms and consequences of bacterial resistance to antimicrobial peptides. Drug Resist Updat. 2016;26:43-57. Epub 2016/05/18. doi: 10.1016/j.drup.2016.04.002. PubMed PMID: 27180309.

31. Snijder J, Reddy VS, May ER, Roos WH, Nemerow GR, Wuite GJ. Integrin and defensin modulate the mechanical properties of adenovirus. J Virol. 2013;87(5):2756-66. Epub 2012/12/28. doi: 10.1128/JVI.02516-12. PubMed PMID: 23269786; PubMed Central PMCID: PMCPMC3571403.

32. Tartaglia LJ, Badamchi-Zadeh A, Abbink P, Blass E, Aid M, Gebre MS, et al. Alpha-defensin 5 differentially modulates adenovirus vaccine vectors from different serotypes in vivo. PLoS Pathog. 2019;15(12):e1008180. Epub 2019/12/17. doi: 10.1371/journal.ppat.1008180. PubMed PMID: 31841560; PubMed Central PMCID: PMCPMC6936886.

33. Waddington SN, McVey JH, Bhella D, Parker AL, Barker K, Atoda H, et al. Adenovirus serotype 5 hexon mediates liver gene transfer. Cell. 2008;132(3):397-409. Epub 2008/02/13. doi: 10.1016/j.cell.2008.01.016. PubMed PMID: 18267072.

34. Kalyuzhniy O, Di Paolo NC, Silvestry M, Hofherr SE, Barry MA, Stewart PL, et al. Adenovirus serotype 5 hexon is critical for virus infection of hepatocytes in vivo. Proc Natl Acad Sci U S A. 2008;105(14):5483-8. Epub 2008/04/09. doi: 10.1073/pnas.0711757105. PubMed PMID: 18391209; PubMed Central PMCID: PMCPMC2291105.

35. Burckhardt CJ, Suomalainen M, Schoenenberger P, Boucke K, Hemmi S, Greber UF. Drifting motions of the adenovirus receptor CAR and immobile integrins initiate virus uncoating and membrane lytic protein exposure. Cell Host Microbe. 2011;10(2):105-17. Epub 2011/08/17. doi: 10.1016/j.chom.2011.07.006. PubMed PMID: 21843868.

36. Kundhavai Natchiar S, Venkataraman S, Mullen TM, Nemerow GR, Reddy VS. Revised Crystal Structure of Human Adenovirus Reveals the Limits on Protein IX Quasi-Equivalence and on Analyzing Large Macromolecular Complexes. J Mol Biol. 2018;430(21):4132-41. Epub 2018/08/20. doi: 10.1016/j.jmb.2018.08.011. PubMed PMID: 30121295.

37. Yu X, Veesler D, Campbell MG, Barry ME, Asturias FJ, Barry MA, et al. Cryo-EM structure of human adenovirus D26 reveals the conservation of structural organization among human adenoviruses. Sci Adv. 2017;3(5):e1602670. Epub 2017/05/17. doi: 10.1126/sciadv.1602670. PubMed PMID: 28508067; PubMed Central PMCID: PMCPMC5425241.

38. Warming S, Costantino N, Court DL, Jenkins NA, Copeland NG. Simple and highly efficient BAC recombineering using galK selection. Nucleic Acids Res. 2005;33(4):e36. Epub 2005/02/26. doi: 10.1093/nar/gni035. PubMed PMID: 15731329; PubMed Central PMCID: PMCPMC549575.

39. Vogt D, Zaver S, Ranjan A, DiMaio T, Gounder AP, Smith JG, et al. STING is dispensable during KSHV infection of primary endothelial cells. Virology. 2020;540:150-9. Epub 2020/01/14. doi: 10.1016/j.virol.2019.11.012. PubMed PMID: 31928996; PubMed Central PMCID: PMCPMC6961814.

40. Cauthen AN, Welton AR, Spindler KR. Construction of mouse adenovirus type 1 mutants. Methods Mol Med. 2007;130:41-59. Epub 2007/04/03. doi: 10.1385/1-59745-166-5:41. PubMed PMID: 17401163.

